# SCHiRM: Single Cell Hierarchical Regression Model to detect dependencies in read count data

**DOI:** 10.1101/335695

**Authors:** Jukka Intosalmi, Henrik Mannerström, Saara Hiltunen, Harri Lähdesmäki

## Abstract

**Motivation:** Modern single cell RNA sequencing (scRNA-seq) technologies have made it possible to measure the RNA content of individual cells. The scRNA-seq data provide us with detailed information about the cellular states but, despite several pioneering efforts, it remains an open research question how regulatory networks could be inferred from these noisy discrete read count data.

**Results:** Here, we introduce a hierarchical regression model which is designed for detecting dependencies in scRNA-seq and other count data. We model count data using the Poisson-log normal distribution and, by means of our hierarchical formulation, detect the dependencies between genes using linear regression model for the latent, cell-specific gene expression rate parameters. The hierarchical formulation allows us to model count data without artificial data transformations and makes it possible to incorporate normalization information directly into the latent layer of the model. We test the proposed approach using both simulated and experimental data. Our results show that the proposed approach performs better than standard regression techniques in parameter inference task as well as in variable selection task.

**Availability:** An implementation of the method is available at https://github.com/jeintos/SCHiRM.

## 1 Introduction

Gene regulatory network (GRN) inference has been a vivid line of research for years. For bulk expression data there exists a wide variety of methods to infer gene regulatory interactions (Marbach *et al.*, 2012). Recently developed single cell measurement technologies provide rich information about the expression levels of different cellular components in individual cells and there is a great promise that these novel technologies could also result in more detailed predictions about the gene regulatory interactions. Along with novel single cell measurement technologies, also new computational techniques for GRN inference have begun to emerge (for recent reviews, see Babtie *et al.*, 2017; Fiers *et al.*, 2018). Many of the proposed GRN inference methods make use of time (or pseudotime) alignment of cells (see e.g. Specht and Li, 2017; Sanchez-Castillo *et al.*, 2018; Papili Gao *et al.*, 2018) and only few approaches can be used to detect dependencies from static data using regression modeling (see e.g. Huynh-Thu *et al.*, 2010; Aibar *et al.*, 2017). Although single cell data provides detailed view on cellular states, the related data analysis tasks remain challenging due to the notable amount of biological and technical variability in single cell data.

Single cell RNA sequencing (scRNA-seq) is one of the most popular experimental techniques to measure gene expression levels at the single cell level (Kolodziejczyk *et al.*, 2015). In general, scRNA-seq data consist of discrete read count measurements, each read count reflecting the expression level of a specific gene in a single cell (Fig. 1 (a)). These data are often described using a Poisson model which is extended to account for overdispersion effects that can originate from various technical and biological sources (Vallejos *et al.*, 2015; Zappia *et al.*, 2017). Thus, the true gene expression patterns in scRNA-seq data can be seen to be covered with overdispersed Poisson noise and the discrete noise needs to be explained away if we wish to make any inference about the underlying signals. The observed read counts can also be affected by other factors, such as the cell sizes, and these factors need to be taken into account when designing statistical models. Hierarchical modeling approaches provide excellent premises to incorporate all these features into a single modeling framework (cf. Vallejos *et al.*, 2015).

**Figure 1:**
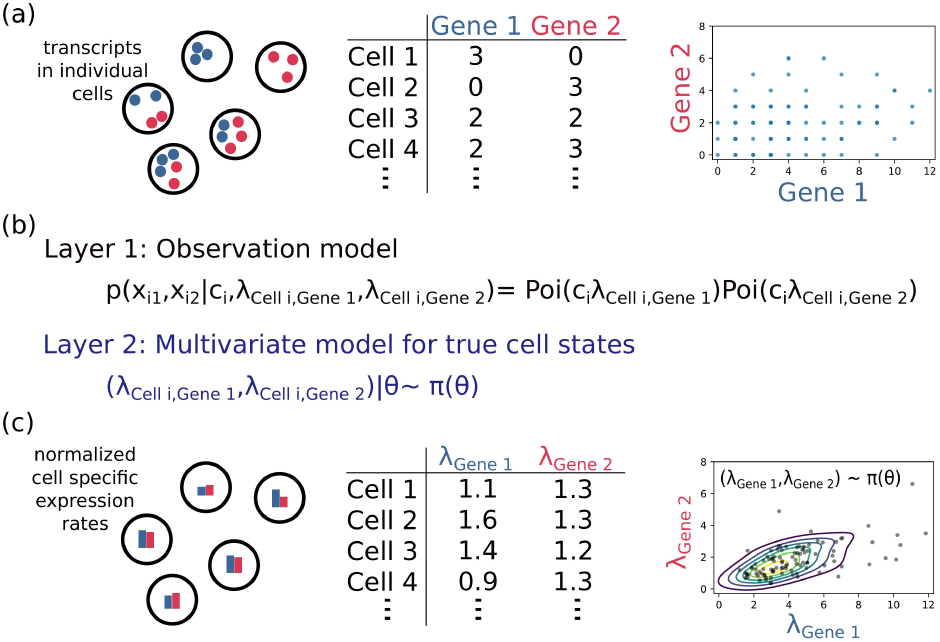
Illustration of scRNA-seq data and the hierarchical modeling approach. (a) The discrete read count data can be collected in a table where columns consists the measured values over all cells. (b) Each count value is assumed to follow the Poisson distribution which is parameterized by the corresponding latent expression level parameter λ _Cell*i*_,_Gene*j*_. The expression level parameters are scaled by normalization constant *c*_*i*_ to take cell size effect into account. The latent expression parameters follow a joint distribution π which contains all information about complex relationships between different genes.(c) The hierarchical approach makes it possible to model the expression levels in a domain where the discrete Poisson type of noise is explained away and, consequently, the underlying dependencies between the genes can be inferred reliably.

In a hierarchical formulation, the stochastic effects in scRNA-seq data can be naturally divided into two layers. These two layers are illustrated in Fig. 1 (b) in a setting in which two genes are observed. The first layer describes the observational uncertainty for discrete read count measurements. The second layer describes the underlying true cellular states which are inherently stochastic due to the stochastic nature of transcriptional processes and other random effects affecting the cell states. The true expression levels are assumed to follow a multivariate distribution (Fig. 1 (c)) which is informative about the possible dependencies between genes and, in the context of GRN inference, we aim to recover these dependencies. This abstraction of the mechanistic grounds of gene expression provides us with a very flexible modeling framework. For instance, normalization information, such as cell size related effects, can be directly incorporated into the observation model without interfering the latent domain where the crucial dependencies are modelled (in the illustration, this is done by multiplying the expression level by *c*_*i*_ in Fig. 1). Further, because we model discrete count data using an appropriate discrete distribution, we do not need any cumbersome and arbitrary data transformations; contrary to standard regression methods that have been proposed in literature.

Modeling scRNA-seq data using the hierarchical formulation (Fig. 1 (b) and (c)) in its most general form is in principle possible but, in practice, this would result in an overly flexible model. Consequently, we need to constrain the presented abstraction in a feasible way to obtain practically useful model which has intuitive parameterization. In this study, we formulate a single cell hierarchical regression model (SCHiRM) which provides us with one possible distribution family and parameterization of the assumed data generating process. To derive SCHiRM, we model the joint distribution of the logarithmic expression rates by conditioning the expression level of a given target gene by the expression levels of candidate input genes. This formulation results naturally in a regression model. Importantly, the SCHiRM accounts for uncertainty in both input and output variables and can be extended in many ways due to its modular design.

We implement the SCHiRM for scRNA-seq data in Python using the probabilistic programming language Stan (Carpenter *et al.*, 2017) which enables easy testing of different variants of the model as well as robust and reliable, sampling based, posterior analysis. Our computational experiments with simulated and real experimental data show that the SCHiRM performs notably better than standard regression techniques.

## 2 Methods

### 2.1 Data

We test the SCHiRM using simulated as well as experimental scRNA-seq data. The experimental data for 239 human K562 erythroleukemia lymphoblast cell line cells is obtained from Klein *et al.* (2015). The counts in the data represent the actual molecule numbers which are obtained by using unique molecular identifier correction (Islam *et al.*, 2013). For details about the data see Klein *et al.* (2015) (see also Supplementary Information, SFig. 1).

### 2.2 Poisson-log normal distribution

We model scRNA-seq data using the Poisson-log-normal distribution which can be defined as a mixture of Poisson distribution with log-normally distributed rate parameter λ (Aitchison and Ho, 1989). Formally, a univariate random variable *x* follows the Poisson-log normal distribution if

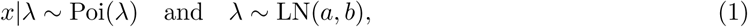

where Poi(λ) is the Poisson distribution with the parameter λ and LN(*a, b*) is the log-normal distribution with parameters *a* and *b*. Further, the mean and variance of *x* depend solely on the mean and variance of λ as

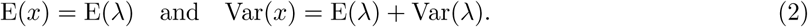

We denote E(λ) = μ and Var(λ) = *σ*^2^. Based on the properties of the log-normal distribution LN(*a, b*), we can write

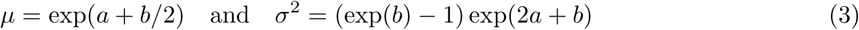

and by solving the equations for *a* and *b*, we can obtain a reparameterized version of the log-normal distribution LN*\**(μ, *σ*^2^). In other words, *Z ∼* LN*\**(μ, *σ*^2^) is equal to

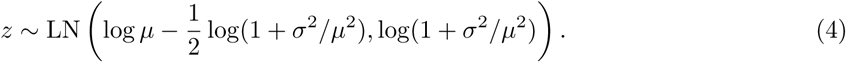

The reparameterization is very convenient in the context of Poisson-log normal distribution because the parameters μ and *σ*^2^ have now a direct connection to the properties of the random variable *x* through the relationships shown in Eq. 2. The mean parameter μ fully determines the mean of *x* and variance parameter *σ*^2^ controls the amount of the overdispersion with respect to the standard Poisson model which is obtained if *σ*^2^ → 0.

### 2.3 Overdispersion

The Poisson-log normal distribution provides flexible means to model discrete read count data and, importantly, this distribution also accounts for overdispersion. In our application, we assume that the amount of overdispersion depends on the mean expression level. To model this dependency quantitatively, we introduce a parametric dispersion function *f >* 0 and assume that

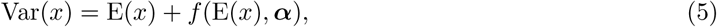

where α includes the dispersion function parameters. Here, the random variable *x* represents the expression level of a single gene. We assume that the functional form of the dispersion function *f* is specified by the modeler. The dispersion function parameters can be fixed based on prior information or learned during the inference as we do in this study.

### 2.4 Incorporating the overdispersion function into the Poisson-log normal model

The overdispersion in the Poisson-log normal model is fully determined by the variance parameter *σ*^2^ of the underlying log-normal distribution LN^*\**^(μ, *σ*^2^). Consequently, *σ*^2^ is the parameter which we wish to link with the dispersion function *f.* This can be done by combining the expressions for Var(*x*) in Eqs. 2 and 5, i.e.

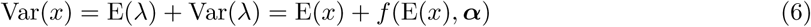

which simply yields that Var(λ) = *σ*^2^ = *f* (μ, α). Now, we can write the model for *x* in the form

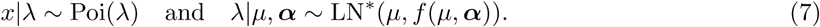

This is a Poisson-log normal model which is parameterized by μ and α given the dispersion function *f.*

### 2.5 Hierarchical regression model

Let us consider *M* input genes in *N* cells and denote the random variables representing their expression levels by *x*_*ij*_ with *i* = 1, *…, N* and *j* = 1, *…, M.* It would be natural to assume that each *x*_*ij*_ follows the Poisson distribution with some (latent) rate parameter *λ*_*ij*_. However, this formulation is infeasible because a fixed expression rate in two different cells may result in totally different expression levels due to the difference in the cell sizes (see e.g. Vallejos *et al.*, 2015, for an illustration). Consequently, the Poisson rate constants need to be scaled according to the cell sizes (and possibly by other factors). For this purpose, we define the cell specific normalization constants

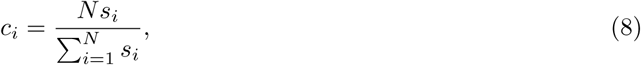

where *s*_*i*_, *i* = 1, *…, N* are the relative cell sizes which can be estimated based on the total mRNA content of the cells (for details, see Subsection 2.6).

Now, given the normalization constants *c*_*i*_, *i* = 1, *…, N* and a dispersion function *f* (μ, *α*), we can set Poisson-log normal distributions for the input genes, i.e.

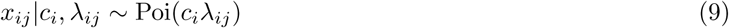

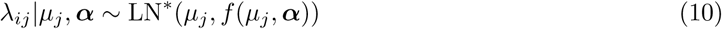

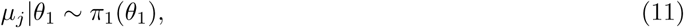

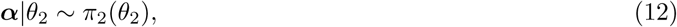

where *λ*_*ij*_ are Poisson rate parameters corresponding to each *x*_*ij*_, *µ*_*j*_ are gene specific mean expression levels, α contains dispersion function parameters, and *π*_1_ and *π*_2_ are the prior distributions for *µ*_*j*_ and α, respectively. Hyperparameters *θ*_1_ and *θ*_2_ are taken to be fixed.

As explained above, our aim is to predict the relationships between a target gene and given candidate input genes indexed by *j* = 1, *…, J.* We denote the random variables representing the expression levels of the target gene in *N* cells by *y*_*i*_, *i* = 1, *…, N* and define the relationship between the inputs and target by

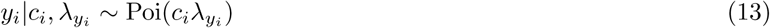

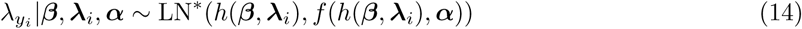

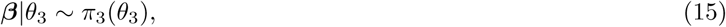

where β = (*β*_0_, *β*_1_, *…, β_M_)* are the regression parameters, ***λ****i* = (*λi*_1_, *λ*_*i*2_, *…, λ*_*iM*_) are the true (latent) expression rate parameters of the input genes,

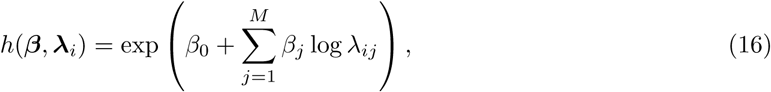

and *π*_3_ is a prior distribution for β. Hyperparameters *θ*_3_ are taken to be fixed. Here, Eqs. 14 and 16 simply imply that the mean of the target gene parameter *λ*_*y*_*i* is predicted using linear regression in the log-space, this mean value is mapped to the linear scale, and the overdispersion function is used to determine the amount of overdispersion in the conditional distribution.

The interpretation of the model is intuitive. The observed counts are realizations from independent Poisson distributions which are parameterized using latent variables *λi*_*j*_ and *λ*_*y*_*i.* In the hierarchical formulation, the latent parameters represent the actual cell specific gene expression levels and, thus, it is natural to formulate also the regression model in this domain. This is analogous to the standard regression analysis which is carried out for log-transformed read count data. However, the crucial difference between the approaches is that in standard regression modeling the gene expression levels are crudely approximated using a data transformation e.g. log *λ*_*ij*_ *≈* log(1 + *x*_*ij*_) and log *λ*_*y*__*i*_ *≈*log(1 + *y*_*i*_) whereas, in the hierarchical model, the cell specific expression levels are explicitly modeled as latent random variables. Further, applying the normalization to the read counts prior the log-transformation e.g. log *λ*_*ij*_ *≈* log(1 + *x*_*ij*_*/c*_*i*_) can bias the analysis. The problems with the log-transformed data become more apparent when low read counts are considered. This is exemplified in Fig. 2. Importantly, our novel approach and standard approaches have the same interpretation of the regression problem but our hierarchical model takes the uncertainty in the input and output data into account and allows modeling without cumbersome and arbitrary data transformations.

**Figure 2:**
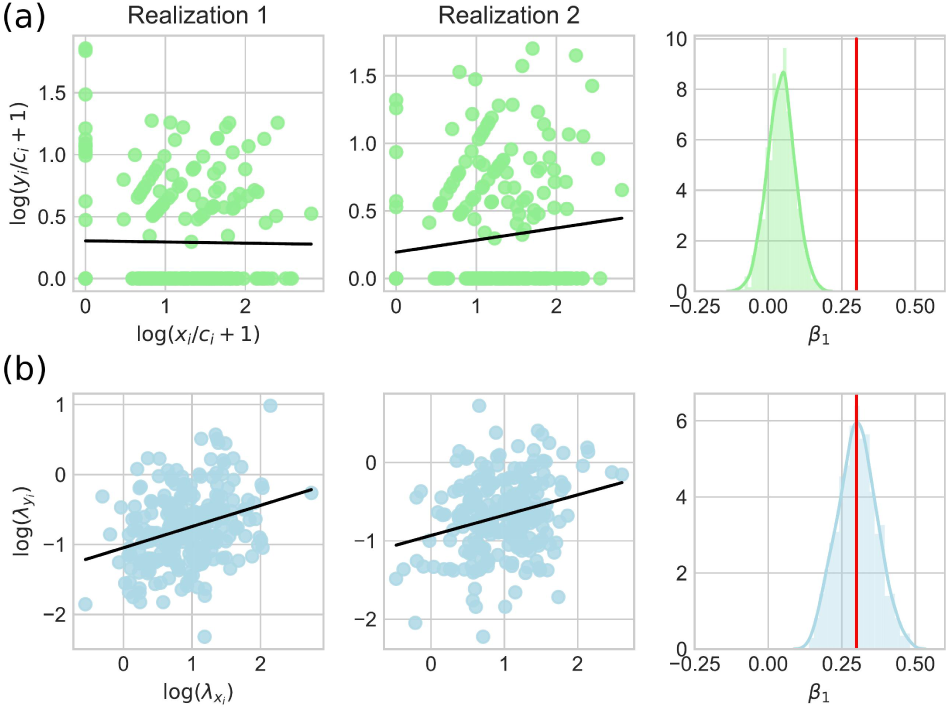
Linear regression for log-transformed normalized counts and latent expression level parameters.1000 data sets (realizations) consisting of read counts for two variables *x* and *y* are simulated from the hierarchical model using *μ*_1_ = 3, *β*β = 1, *β*1 = 0.3, α = 0.3, and normalization constants *c*_*i*_ which are obtained from the real data (Klein *et al.*, 2015). The cell size normalized read counts are transformed into the log-scale and standard linear regression analysis is carried out. The obtained regression lines are shown for two data sets to illustrate how the direct normalization of read counts and logarithmic transform can bias the analysis result. The normalized histogram of estimated values for *β*1 over all data sets further illustrates how the estimates are in general unreliable and often the positive slope cannot be recovered. Together with the histogram, the true value of *β*_1_ is indicated using a gray vertical line. The same as in (a) but now the regression analysis is carried out for the latent parameters in the hierarchical model. In this case, the positive slope can be recovered robustly.

### 2.6 Choice of dispersion function, prior specification and cell size estimation

In general, there are no restrictions how the positive dispersion function *f* can be constructed. In this study, we use the form *f* (μ, α) = *α μ*^2^ which has some desired analytical properties (see Supplementaty Information) and can also be linked with dispersion modeling in the context of bulk RNA-seq data (see e.g. Robinson and Smyth, 2008).

Model parameters do not have prior dependencies and we use proper prior distributions. More precisely, we define

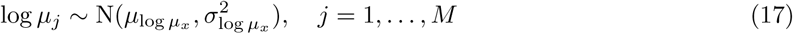

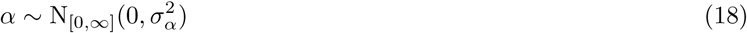

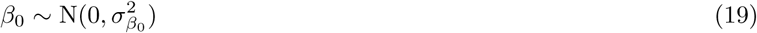

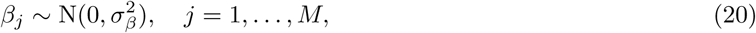

where N(*a, b*) is the normal distribution and N[_*l,u*_](*a, b*) is the truncated normal distribution in the interval [*l, u*] with mean and variance parameters *a* and *b*. The global parameters 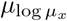 and 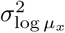 can be calibrated directly from data prior to the analysis (see Supplementary Information, SFig. 1). Throughout this study, we fix *σ*^*α*^ = 0.15 to constrain the dispersion level in a reasonable range. The 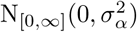 prior for dispersion parameter weights low dispersion levels slightly and, thus, incorporates the prior assumption that the data may be pure Poisson noise. Informative prior for *β*_0_ is hard to form and we have simply assume that 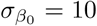. Based on the nature of scRNA-seq data, we would presume that the range of the parameters *β*_*j*_ is roughly from *-*3 to 3, and, consequently, we set 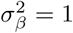 The relative cell sizes *s*_*i*_, *i* = 1, *…, N* are estimated by means of the total expression levels

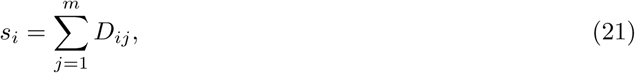

where *D* is the full read count data consisting of *N* cells and *m* genes.

### 2.7 Standard regression approaches

We compare the variable selection performance of SCHiRM with four standard regression approaches: ordianary least squares regression (OLS) (Hastie *et al.*, 2001), least absolute shrinkage and selection operator regression (LASSO) (Tibshirani, 1996), elastic net regression (ENET) (Zou and Hastie, 2005), and ridge regression (RIDGE) (Hastie *et al.*, 2001). In addition, we include Pearson correlation coefficient (PCC) to the comparison. Before applying these methods, we apply the log-transformation to the normalized count data i.e. the input for regression is log(1 + *x*_*ij*_*/c*_*i*_) and output is log(1 + *y*_*i*_*/c*_*i*_). For LASSO, ENET, and RIDGE regression, we select the regularization parameters using cross-validation.

### 2.8 Receiver operator characteristic analysis

We compare the variable selection performance of different methods using receiver operator characteristic (ROC) analysis and area under the curve (AUC) measure (Fawcett, 2006). For this purpose, we need to define the scores which indicate how strongly a certain input variable is linked with the output variable. In the case of standard regression approaches, we simply define score*j* = *|β*_*j*_*|, j* = 1, *…, M,* where *β*_*j*_ are the estimated regression parameters. For PCC, we define score*j* = *r*_*j*_, where *r*^*j*^ is the estimated PCC between the *j*th input and output. For SCHiRM, we define two different scores. The first one is based on probability and takes the form

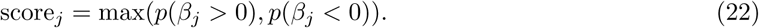

The second one is based on the posterior mean of *β*_*j*_ i.e.

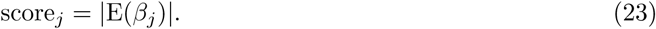

These scores can be easily estimated using the posterior samples and, in the following, the results computed using these scores are denoted by SCHiRMp and SCHiRMpm, respectively.

### 2.9 Computational implementation

We implement the SCHiRM in Python using the probabilistic programming language Stan (Carpenter *et al.*, 2017). Posterior analysis is carried out by means of Markov chain Monte Carlo (MCMC) sampling using the no-U-turn sampler (Hoffman and Gelman, 2014) available in Stan. For all posterior analysis tasks, we sample four independent MCMC chains, each chains consisting of 500 samples. The first half of the samples is discarded as a burn-in and the rest of the samples are used for posterior analysis. The convergence is monitored by means of the potential scale reduction factors (Gelman *et al.*, 2013) and the convergence diagnostics should be checked routinely when using SCHiRM (see also Supplementary Information, SFig. 2). For standard regression approaches, we use the implementations available in the Python package scikit-learn (Pedregosa *et al.*, 2011). For regularized regression models, the regularization parameters are selected using automatized cross validation routines in scikit-learn package (LassoCV, RidgeCV, ElasticNetCV) using default settings.

## 3 Results

### 3.1 An example inference task

Practical application of the SCHiRM is straightforward as almost the whole inference procedure can be automated. In the first step, the full data set is used to estimate the cell-specific normalization constants. The second step is to define the target gene (output) and the potential regulators of the gene (inputs) and run the inference algorithm. After running the algorithm, the user can plot a posterior summary of the model parameters as well as two different scores for inferred regulatory interactions (Figs. 3 and 4).

**Figure 3:**
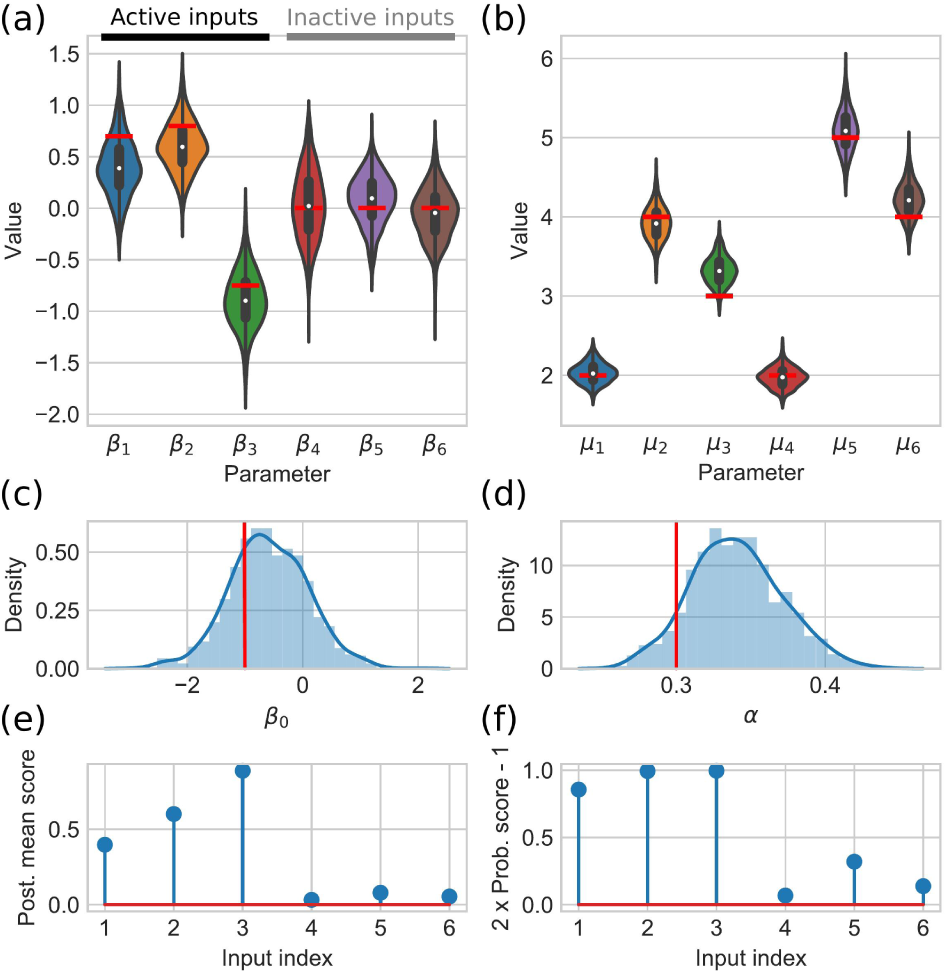
Example of posterior summary and variable selection scores for a simulated data. (a) The estimated posterior distributions for regression coefficients illustrated using violin plots. (b) The estimated posterior distributions for mean parameters of input genes are illustrated using violin plots. (c) The posterior distribution of the intercept is illustrated using normalized histogram and kernel density estimate. (d) The estimated posterior distribution of the dispersion parameter is illustrated using normalized histogram and kernel density estimate. On top of the posterior distributions, the underlying true parameter values are indicated by red bars. (e) and (f) show the obtained posterior mean (Eq. 23) and probability (Eq. 22) based scores, respectively.

**Figure 4:**
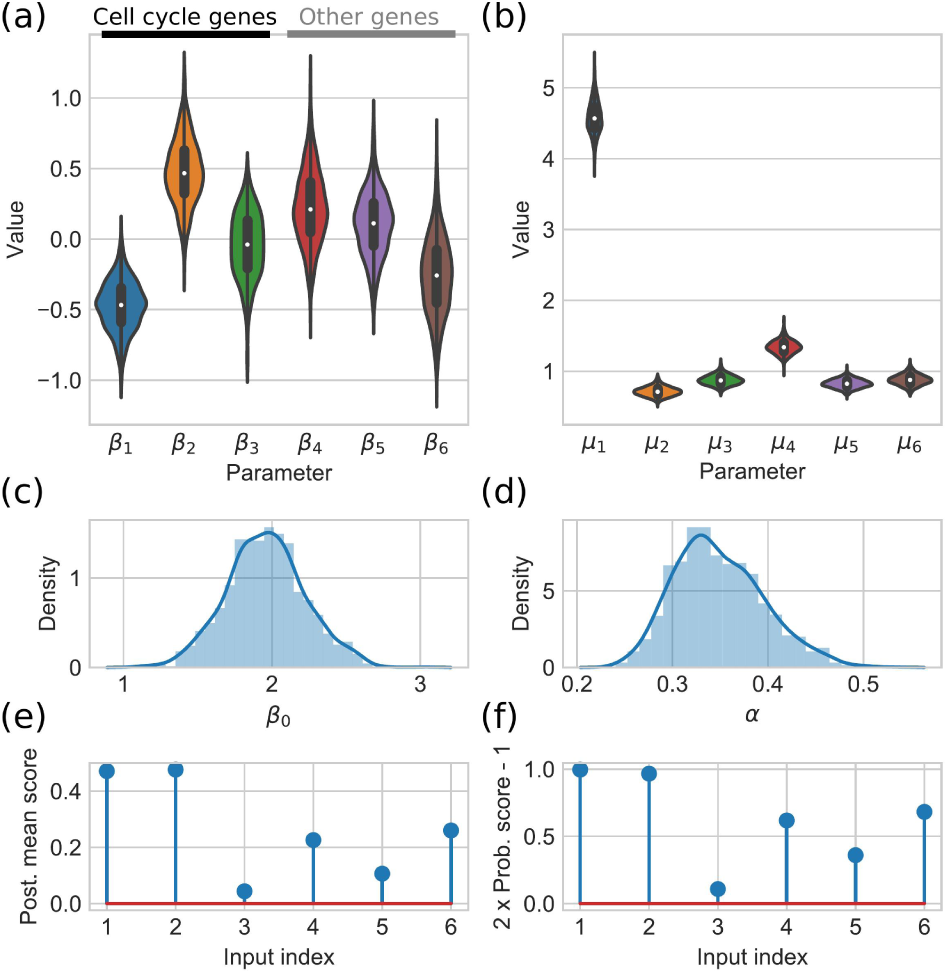
Example of posterior summary for real data (for details about the illustration, see the caption of Fig. 3). In this example inference task, the target gene and first three input genes are related to the cell cycle and rest of the input genes are selected randomly.

To illustrate the inference process, we generate a simulated input data consisting of six genes and the corresponding output gene data assuming that three of the input genes regulate the output gene (for details about data simulation, see Supplementary Information). After running the inference algorithm for these genes, we obtain the summary which is shown in Fig. 3. We see that the posterior distributions for the model parameters are in a good agreement with the underlying true parameters. Further, both, probability and posterior mean based, scores show higher values for the input variables which are affecting the target gene.

A similar illustration can be carried also for real K562 data. In these data, genes which are related to the cell cycle are known to exhibit strong correlations (Klein *et al.*, 2015). In our illustration, we select the target gene to be one of the cell cycle related genes (the list of 38 cell cycle genes are collected from Klein *et al.*, 2015; Whitfield *et al.*, 2002). Further, we define a set of input genes which consists of three other cell cycle related genes and three genes which are not related to cell cycle. Now, the cell cycle related output gene should correlate with the three cell cycle related input genes but not with the other three genes. The summary of the inference is shown in Fig. 4: The scores for two of the cell cycle related inputs are clearly on a higher level than the other scores. The third cell cycle related gene does not seem to correlate with the target gene. The inputs 4, 5, and 6 which are not related to the cell cycle obtain clearly lower scores than strongly correlating cell cycle genes. Further, the posterior distributions of the model parameters are located in reasonable and interpretable ranges.

### 3.2 Variable selection performance with simulated data

One of the most important features of SCHiRM is its variable selection capability based on the scores defined in Section 2.8. We illustrate this capability by simulating data in ten different settings and comparing the variable selection performance of SCHiRM and with standard regression approaches by means of ROC analysis. In different settings, we alter the number of input genes (*M)* and the number of input genes linked with the output gene (*M*_act_) as indicated in the result figures. In all simulations, we have fixed *α* = 0.3 and the truncation range for the input gene mean expression levels in logarithmic scale is [log(0.5), log(100)] (for details about data simulation, see Supplementary Information).

Based on the ROC analysis, the SCHiRM outperforms the standard regression approaches and PCC in all ten settings (Fig. 5 (a)). Interestingly, the performance changes only slightly when the dimension of the input increases. As expected, the SCHiRM performs best in all test cases. Thus, we can presume that the SCHiRM will also perform better for real data given that the assumptions which are made about the data generating process are reasonably realistic.

**Figure 5:**
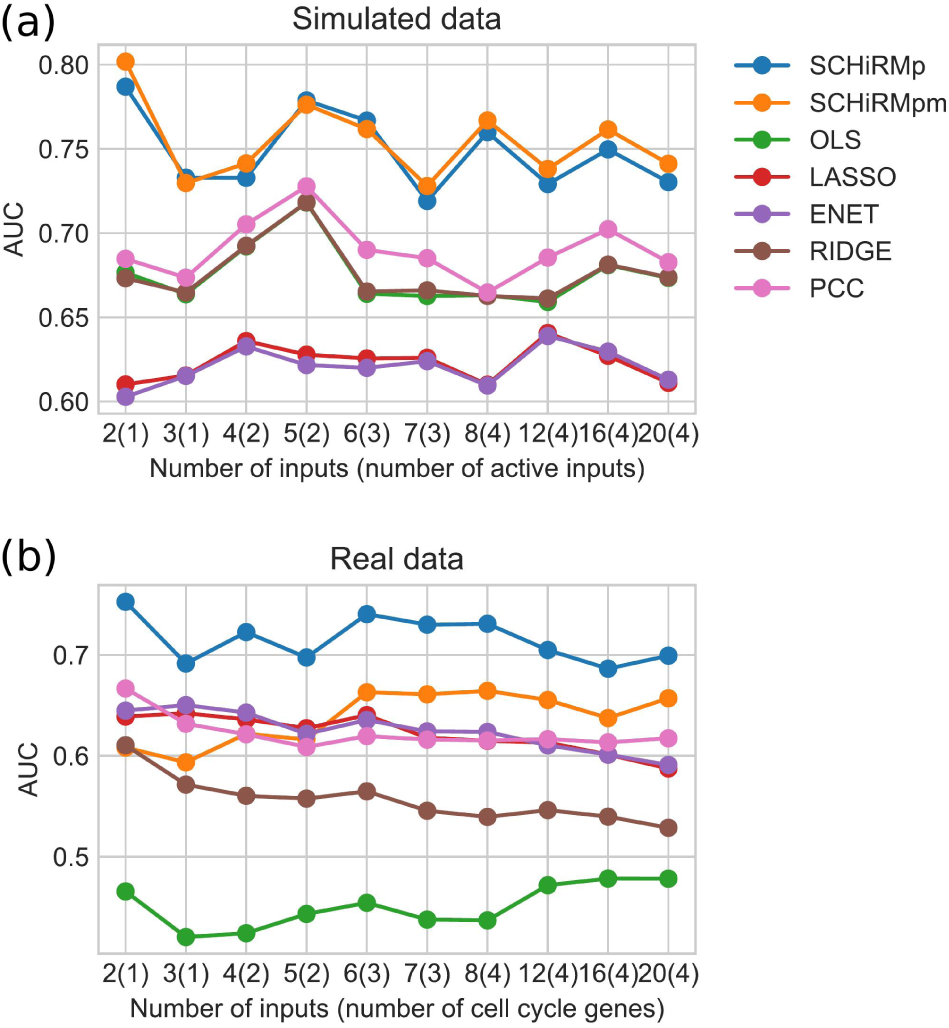
Variable selection performance for simulated and real data. For each number of inputs, the AUC values are computed by considering 100 data sets and carrying out ROC analysis for all *M* × 100 (score, true label) pairs. The number of active inputs which truly affect the expression rate of the target gene are shown in parenthesis. SCHiRMp and SCHiRMpm denote the outcomes which are computed using probabilistic and posterior mean scores for SCHiRM, respectively.

### 3.3 Accuracy of parameter estimates

In addition to variable selection performance we also observed how well the underlying regression model parameters can be estimated by SCHiRM and other methods. Fig. 6 summarizes the parameter estimation performance for all considered methods in all simulated settings by showing the squared error in estimates plotted against the true value. The error in SCHiRM estimates (posterior mean values) for the regression coefficients and intercept is reasonably small over the whole range of considered values. If the true value of regression coefficient is exactly zero or very close to it, the error tends to be larger. However, this occurs rarely enough to produce a too large number of false positives in variable selection context as shown by the ROC analysis above. The standard regression approaches perform poorly in the parameter estimation task, especially if the underlying dependency is strong (Fig. 6). In addition, the parameter estimates provided by the standard techniques tend to be less accurate if the true coefficient or the intercept is negative (Fig. 6). We conclude here that if read count data originates from the hierarchical model, the standard regression techniques applied to log-transformed data can result in severely biased parameter estimates. On the other hand, the SCHiRM performs well in the parameter estimation task. A more detailed summary of the SCHiRM parameter estimates is given in Supplementary Information (SFigs. 3 and 4).

**Figure 6:**
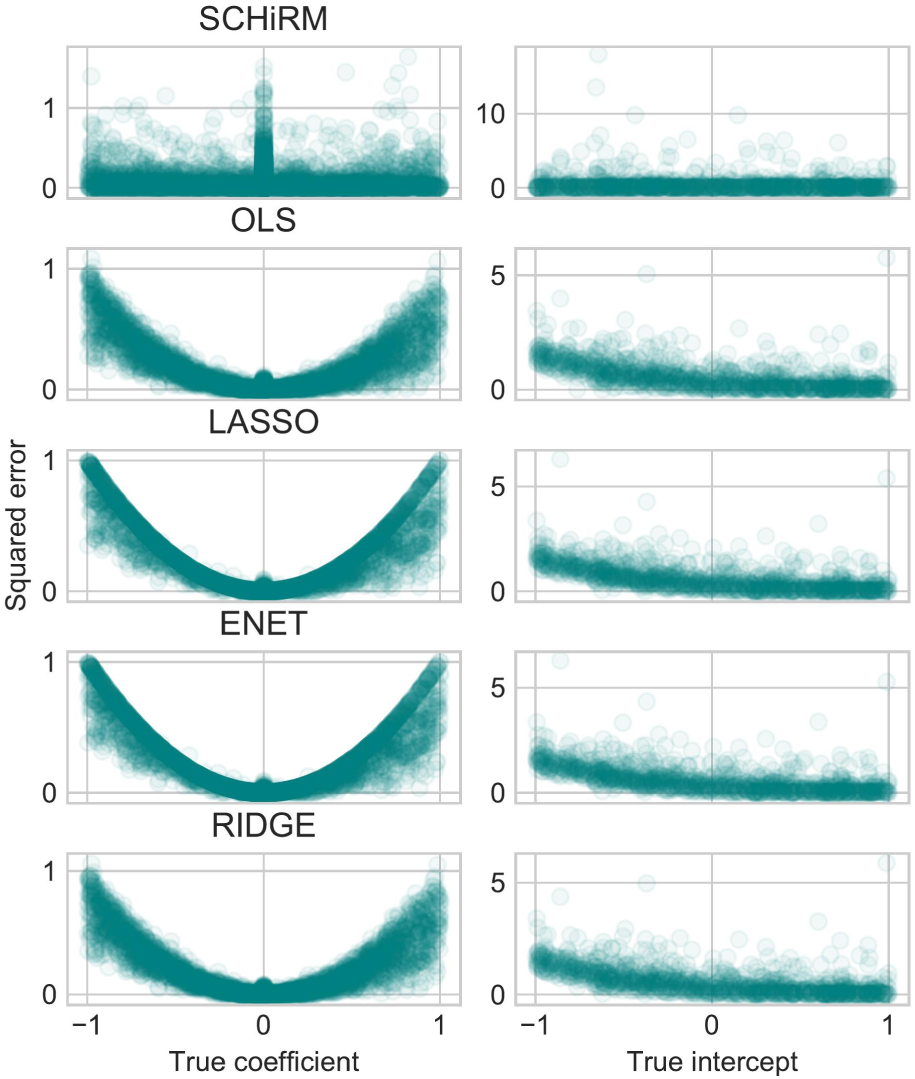
The squared error in parameter estimates plotted against the true underlying parameter value. Each row shows the results for the indicated method. The left panel shows the squared errors for regression coefficients *β*_*j*_ and the right panel for the intercept *β*_0_. The figure shows the squared errors for all parameter estimates obtained from the ten different settings each involving 100 inference tasks.

### 3.4 Variable selection with real data

To test how well the SCHiRM performs in the analysis of real data, we make use of the cell cycle related genes that we used also in the posterior inference example above. In each setting with a fixed dimension of input, we draw a random target (output) gene from the list of cell cycle genes. To construct the input, we further draw random genes form the list of cell cycle genes as well as some number of genes that are not among the cell cycle genes. Now it is reasonable to assume that the cell cycle related target gene will depend more on the cell cycle genes among the inputs than the other input genes. The true and false classes for ROC analysis can thus be formed by assuming that there is true dependency between the cell cycle genes but the dependencies between cell cycle genes and other genes is much weaker or negligible and can be considered to form the false class.

In the context of real data, we use the same input dimension settings as we did with the simulated data. In each setting, we draw 100 distinct input-output pairs, run the inference for these data, and carry out ROC analysis based on the inference results. The result of ROC analysis is summarized in Fig. 5 (b). The results look very similar to what we obtained with simulated data. However, probability based score for SCHiRM seems to be performing better than the posterior mean based score and basic regression modeling (OLS) seems to be performing notably worse than any other approach. We conclude here that also in the case of real data it is beneficial to use the SCHiRM.

## 4 Discussion

In general, statistical treatment of scRNA-seq data is challenging due to the high levels of biological and technical noise (Liu and Trapnell, 2016). Further, co-expression analysis and gene regulatory network inference are among the most challenging data analysis tasks in the context of single cell data (Liu and Trapnell, 2016). Even though there exists GRN inference methods for single cell data, only few studies explore global co-expression patterns by means of regression techniques. When regression techniques are used, they are typically applied directly to (possibly transformed) read count data (see e.g. Dixit *et al.*, 2016; Aibar *et al.*, 2017). In this study, we introduce a principled abstraction to model read count data in a hierarchical manner and build on top of this abstraction to construct the single cell hierarchical regression model (SCHiRM) which can be used to detect regulatory interactions. We exemplify the applicability of the SCHiRM using simulated and experimental data, and our results show that the benefits of using hierarchical formulation are obvious.

In the current implementation of SCHiRM, we assume that the samples originate from one cell type and that we observe the actual molecule numbers which can be obtained e.g. by using unique molecular identifiers (UMIs) (Islam *et al.*, 2013). In this setting, the cell-to-cell variability in the sample acts as a natural perturbation and provides information about co-expression patterns (see e.g. Padovan-Merhar and Raj, 2013) which are then captured by means of regression modeling. As we have shown in this study, the SCHiRM provides a powerful model class within this setting and, given the generality of our modeling approach, the model can be easily extended to be applicable also for more versatile data sets. For instance, if Perturb-seq data (Dixit *et al.*, 2016) is available, the known perturbation effects can be incorporated into the regression model with some uncertainty about the effectiveness of the perturbation. Extended versions of SCHiRM can also be contructred to study multiomics data (Kelsey *et al.*, 2017) in an integrative manner.

The statistical power of SCHiRM does not come without cost. The inference can be run robustly without numerical problems, but due to the hierarchical structure of the model, the posterior sampling can become computationally heavy if a large number of regressor genes are included in the analysis. Although genome-wide network inference is rarely a desired mode of analysis, straightforward application of SCHiRM to genome wide studies is currently out of reach. One way to scale up the method would be to design an appropriately regularized version of the model with an efficient computational strategy to tune the regularization parameters, and then simply use optimization techniques to fit the model. Besides the regularization approach, it could be possible to scale up the method by deriving a variational inference approximation for the posterior. We regard especially the variational inference approach an exiting direction for future development of the SCHiRM.

## 5 Conclusions

We introduce a novel approach to model single cell read count data in a hierarchical manner and provide a computational implementation which can be easily applied to detect dependencies in scRNA-seq data. We implement the method in Python using the probabilistic programming language Stan (Carpenter *et al.*, 2017) which makes our approach easily extendable and applicable in versatile applications. Our computational experiments with simulated and experimental data show that our approach outperforms standard regression techniques in both variable selection and parameters estimation tasks.

## Acknowledgements

The authors acknowledge the computational resources provided by the Aalto Science-IT project.

## Supplementary Information

### 1 **Model formulation with the dispersion function** *f (μ, α)* = *α μ*^2^

If *f* (μ, α) = *αμ*^2^, our custom parameterized Poisson-log normal distribution takes the form

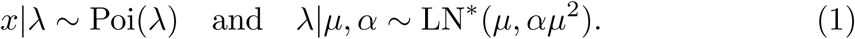

An equivalent formulation for the distribution of λ (or log λ) can be obtained by considering the alternative parameterization in Eq. 4 and by simplifying the formulas which results in

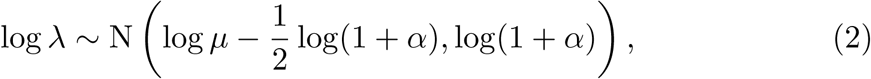

where N(*a, b*) is the normal distribution with the mean *a* and variance *b*.

Given these properties, it is convenient to rewrite the Poisson-log normal distributions for the input genes and the target gene in the form

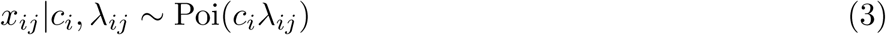

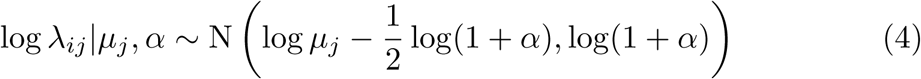

And

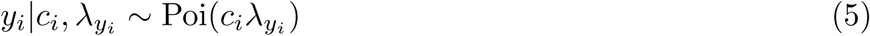

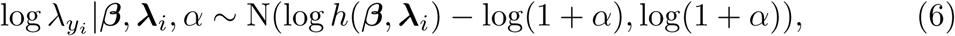

Where

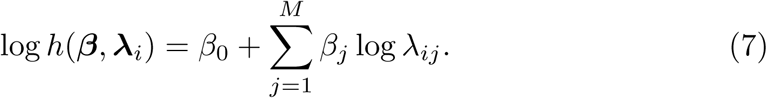

### 2 Simulated data

We simulate artificial data based on statistical characteristics which are extracted from the real K562 data set. First, we estimate the relative cell sizes *s*_*i, i*_ = 1, *…, N* = 239 based on the total mRNA content of the cells and compute the corresponding normalization constants *c*_*i*_. Second, we compute the mean read count levels for all genes and fit a normal distribution to these data in logarithmic scale (at this point, genes which are not expressed at least in one cell are excluded). The fitted normal distribution shows that the data contains a notable amount of lowly expressed genes (SFig. 1) and to maintain a reasonable signal to noise ratio in simulated data, we use a truncated version of the normal distribution to sample the values for log *μ*_*j*_ Our data simulation procedure is outlined below.

In our computational experiments, we consider simulated data pairs (*X, y*) where *X* is a *N* ×*M* matrix containing the read counts of input genes and *y* is *N* -element vector containing the read counts of the target gene. A single pair *(X, y*) where *M*^act ≤^*M* input genes are linked to the output can be simulated using the following procedure:

1. Draw the mean expression rates for input genes as log *u*_*j*_ ∼ N[_*l,u*_](*a, b*), *j* = 1, *…, M,* where *a* and *b* are estimated form an existing data set and [*l, u*] indicates the truncation range.
2. Simulate cell-specific expression rates and randomly draw read counts for the input genes

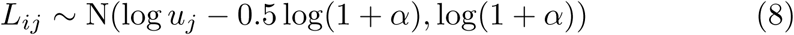

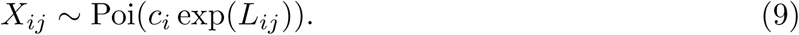
3. Simulate regression coefficients for active and inactive input genes as *β*_*j*_ ∼ *U* (*-*1, 1), *j* = 0…, *M*_act_ and set *β*_*j*_ = 0 for *j* = *M*_act_ + 1, *…, M.*
4. For *i* = 1, *…, N,* simulate stochastic, cell-specific expression rates and randomly draw read counts for the target gene

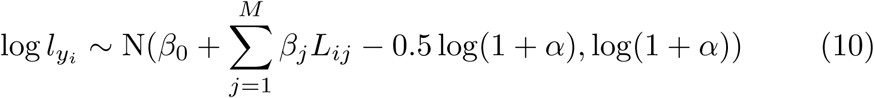

In practice, the exact normalization constants are never available and they need to be estimated from data. To generate realistically blurred estimates of the true normalization constants for computational experiments with simulated data, we generate a data set of 5000 genes through the steps 1 and 2 in the above procedure and estimate the normalization constants form these data. These estimated normalization constants are then used in all experiments with simulated data (SFig. 1).

### 3 Supplementary figures

**SFig. 1.**
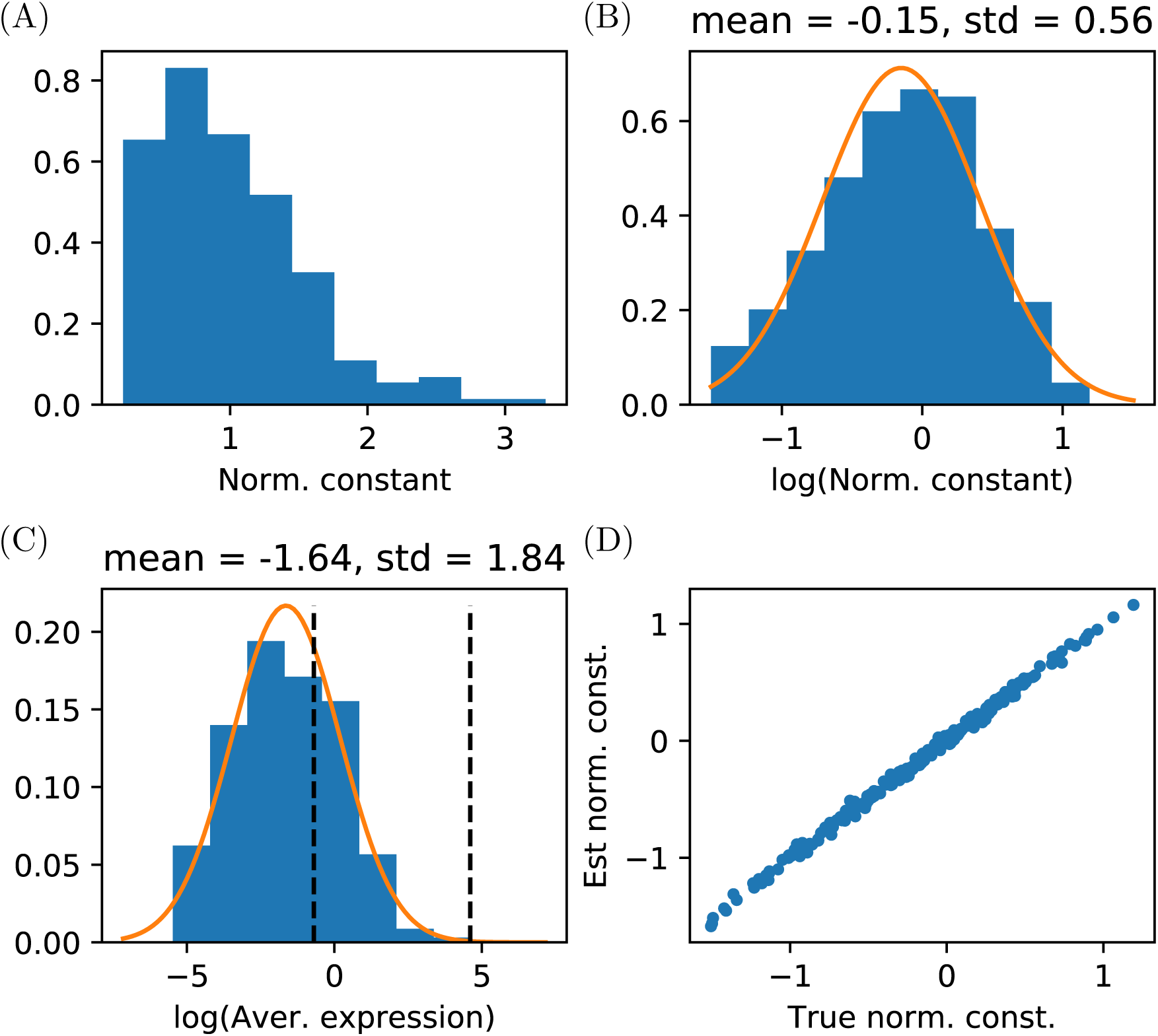
Features computed from the human K562 cell line data set. (A) Histogram of the cell specific normalization constants. (B) Histogram of the logarithmic cell specific normalization constants and the normal distribution which is fitted to these data. The estimated parameters for the normal distribution are given on top of the figure. (C) Histogram of the logarithmic average expression levels over all genes and the normal distribution which is fitted to these data. The estimated parameters of the normal distribution are given on top of the figure. The vertical dashed lines show the truncation range [log(0.5), log(100)] which is used when artificial data is simulated. (D) True logarithmic normalization constants (originating from real data) plotted against the logarithmic normalization constants which are estimated from simulated data. The estimated constants agree reasonably well with the true constants but small jitter can be observed.

**SFig. 2.**
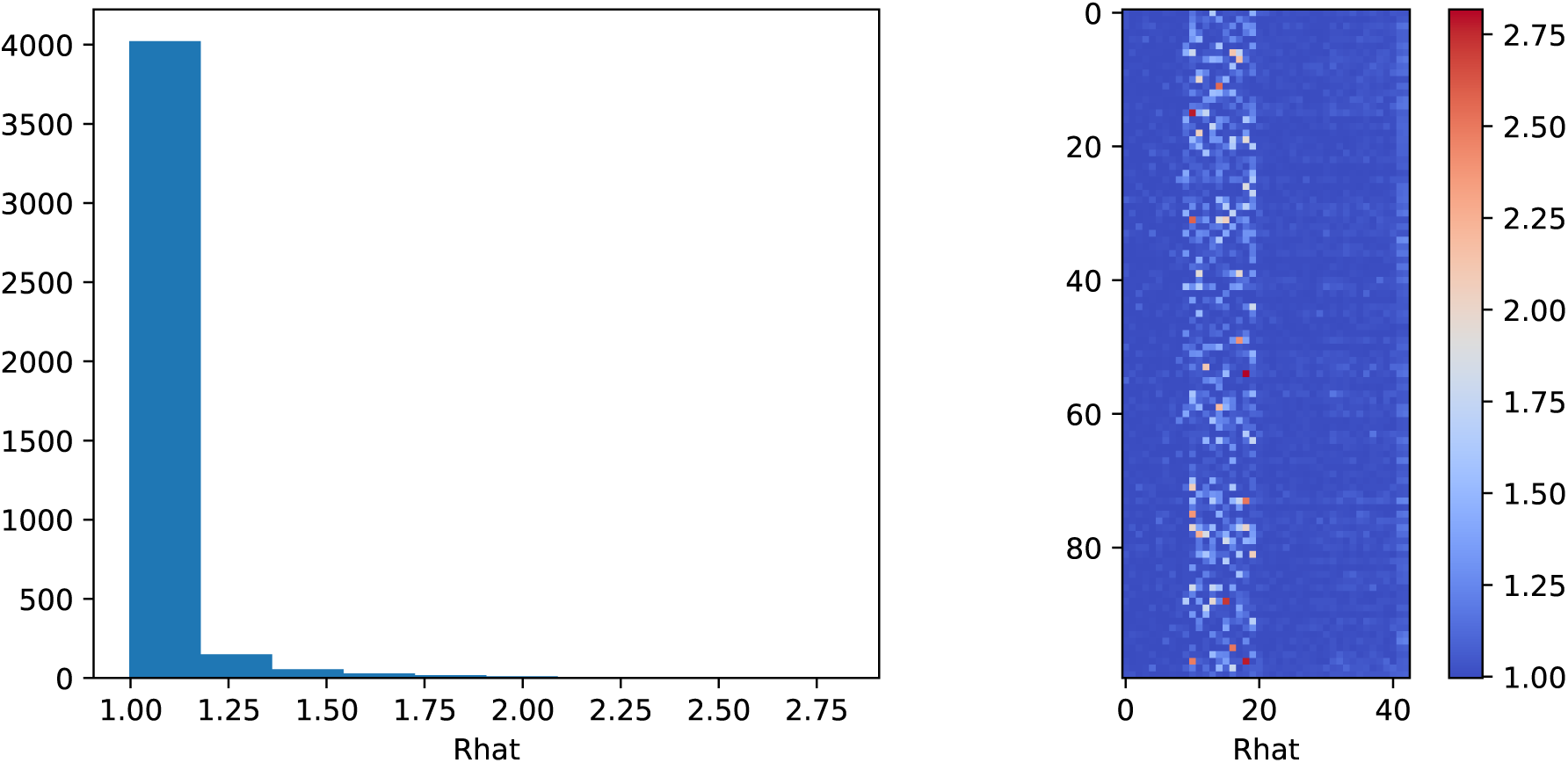
Summary of convergence diagnostics for the analysis of 100 randomized real data sets with *M* = 20 and *M*_act_ = 4 (see Main text, Results Section for details how real data is used here). The potential scale reduction factors (Rhat) are computed for all SCHiRM parameters except for *λ*_*ij*_ and *λ*_*yi*_ which typically do not cause convergence problems. The histogram shows that only few potential scale reduction factors have values greater than 1.1. The heat map shows the computed potential scale reduction factors for individual inference runs (rows) for all parameters of interest (columns). Bad convergence performance may occur if the expression level of one or more inputs is very low and the corresponding regression coefficient becomes unidentifiable. In practice, too lowly expressed genes can often be removed from the data prior to the analysis but the convergence diagnostics should still be always checked routinely.

**SFig. 3.**
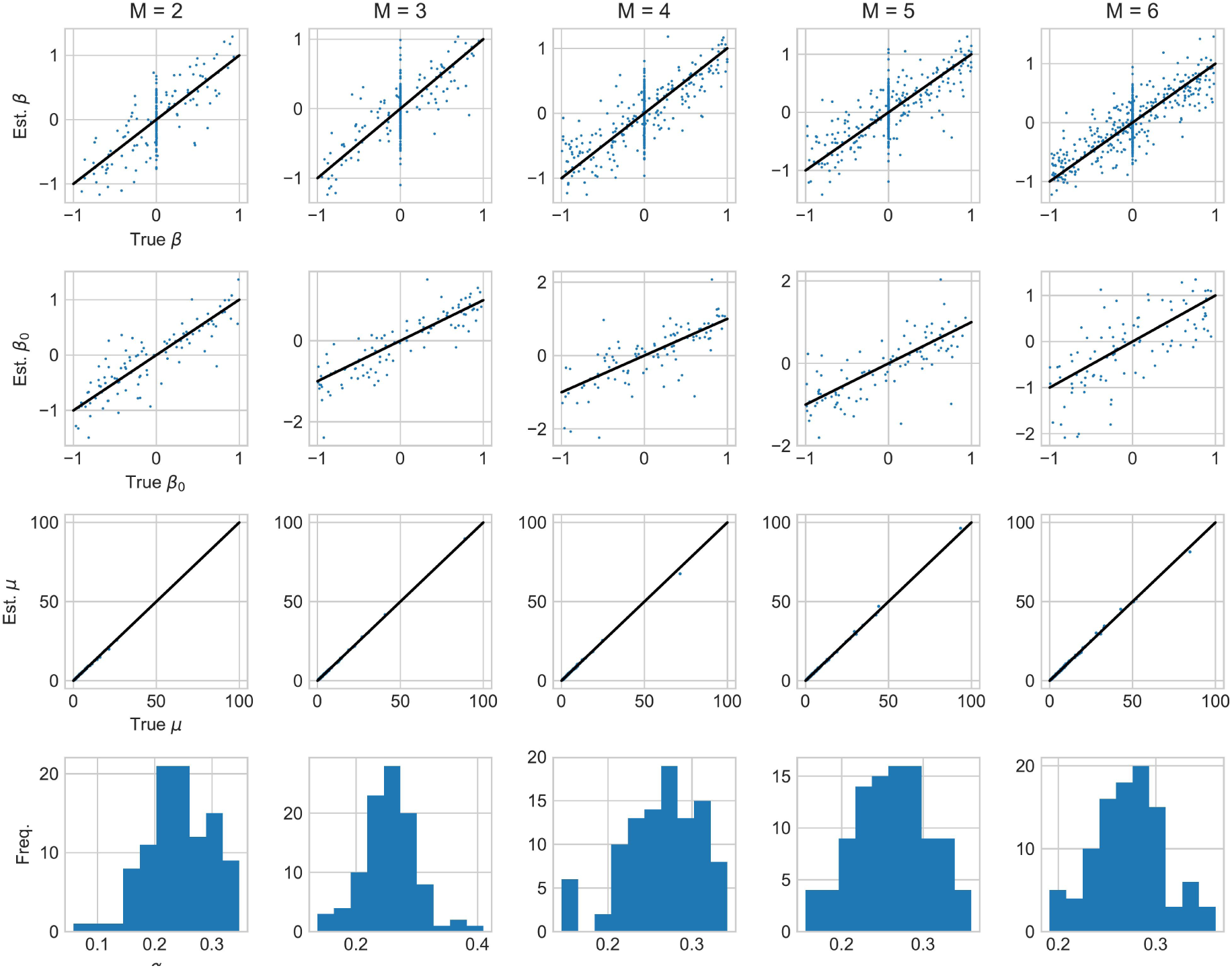
Illustration of SCHiRM parameter estimation performance with varying input dimension (*M* = 2, *…,* 6). For regression coefficients *β*_*j*_, intercept *β*_0_, and mean input levels *μ*_*j*_, the performance is illustrated by plotting the true parameter values against the estimated posterior mean values. The lower panel shows the histograms for the estimated posterior means of dispersion parameter α (in the simulations, *α*_true_ = 0.3). The shown parameter estimates are obtained by running the inference for 100 simulated data sets for each shown input dimension.

**SFig. 4.**
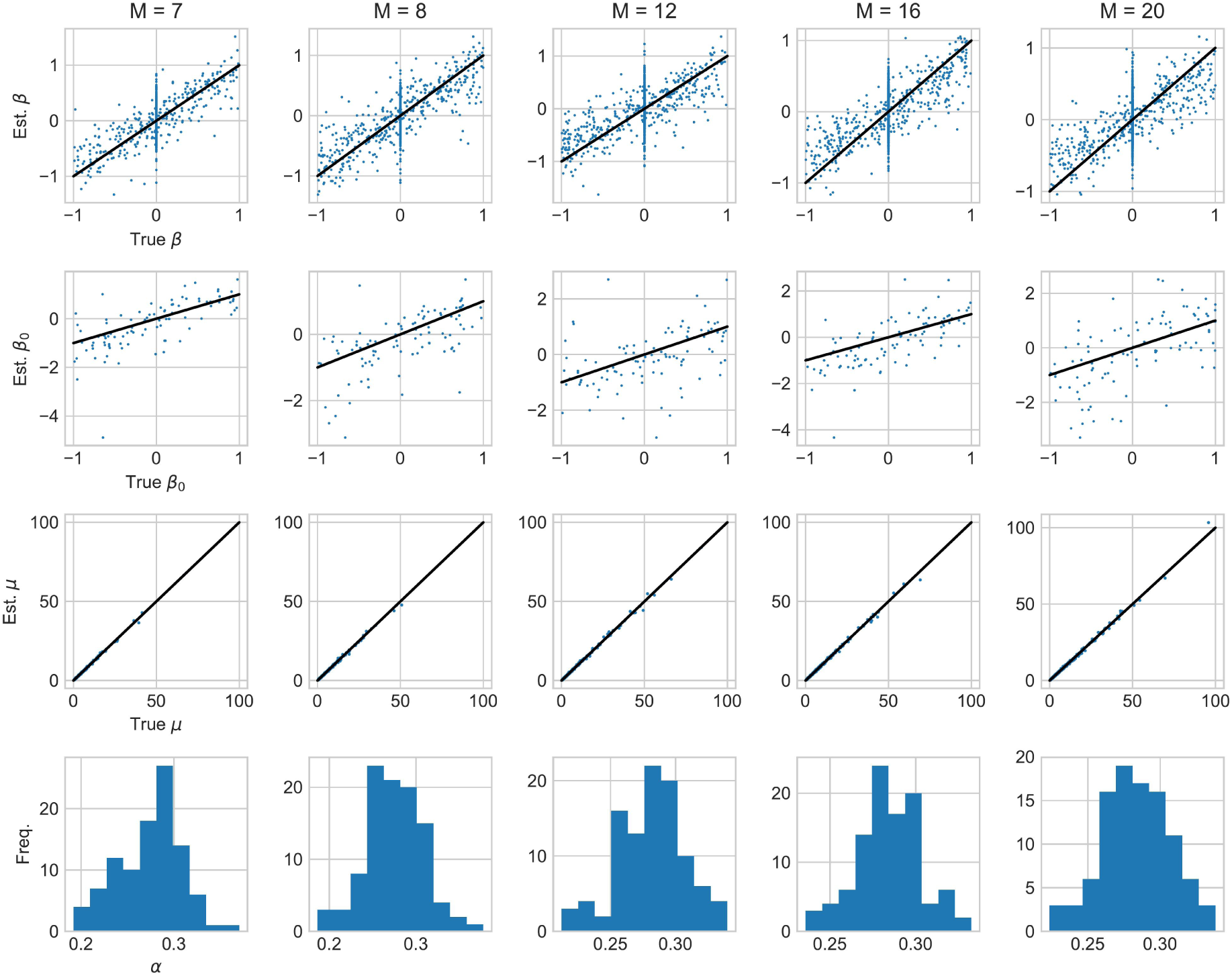
Illustration of SCHiRM parameter estimation performance with varying input dimension (*M* = 7, 8, 12, 16, 20). For regression coefficients *β*_*j*_, intercept *β*_0_, and mean input levels *μ*_*j*_, the performance is illustrated by plotting the true parameter values against the estimated posterior mean values. The lower panel shows the histograms for the estimated posterior means of dispersion parameter α (in the simulations, *α*_true_ = 0.3). The shown parameter estimates are obtained by running the inference for 100 simulated data sets for each shown input dimension.

## References

Aibar, S. et al. (2017). SCENIC: single-cell regulatory network inference and clustering. Nature methods, 14(11), 1083–1086.

Aitchison, J. and Ho, C. H. (1989). The Multivariate Poisson-Log Normal Distribution. Biometrika, 76(4), 643–653.

Babtie, A. et al. (2017). Learning regulatory models for cell development from single cell transcriptomic data. Current Opinion in Systems Biology, 5, 72–81.

Carpenter, B. et al. (2017). Stan: A Probabilistic Programming Language. Journal of Statistical Software, 76(1).

Dixit, A. et al. (2016). Perturb-Seq: Dissecting Molecular Circuits with Scalable Single-Cell RNA Profiling of Pooled Genetic Screens. Cell, 167(7), 1853–1866.e17.

Fawcett, T. (2006). An introduction to ROC analysis. Pattern Recogn Lett, 27(8), 861–874.

Fiers, M. et al. (2018). Mapping gene regulatory networks from single-cell omics data. Briefings in Functional Genomics, page elx046.

Gelman, A. et al. (2013). Bayesian Data Analysis. Chapman & Hall/CRC Texts in Statistical Science, 3rd edition.

Hastie, T. et al. (2001). The Elements of Statistical Learning. Springer New York Inc., New York, NY, USA.

Hoffman, M. D. and Gelman, A. (2014). The No-U-turn sampler: adaptively setting path lengths in Hamiltonian Monte Carlo. Journal of Machine Learning Research, 15(1), 1593–1623.

Huynh-Thu, V. A. et al. (2010). Inferring regulatory networks from expression data using tree-based methods. PLOS ONE, 5(9), 1–10.

aIslam, S. et al. (2013). Quantitative single-cell RNA-seq with unique molecular identifiers. Nature Methods, 11(2), 163–166.

Kelsey, G., Stegle, O., and Reik, W. (2017). Single-cell epigenomics: Recording the past and predicting the future. Science, 358(6359), 69–75.

Klein, A. M. et al. (2015). Droplet Barcoding for Single-Cell Transcriptomics Applied to Embryonic Stem Cells. Cell, 161(5), 1187–1201.

Kolodziejczyk, A. A. et al. (2015). The Technology and Biology of Single-Cell RNA Sequencing. Molecular Cell, 58(4), 610–620.

Liu, S. and Trapnell, C. (2016). Single-cell transcriptome sequencing: recent advances and remaining challenges [version 1; referees: 2 approved]. F1000Research, 5(182).

Marbach, D. et al. (2012). Wisdom of crowds for robust gene network inference. Nat Methods, 9(8), 796–804.

Padovan-Merhar, O. and Raj, A. (2013). Using variability in gene expression as a tool for studying gene regulation. Wiley Interdiscip Rev Syst Biol Med, 5, 751–759.

Papili Gao, N. et al. (2018). Sincerities: inferring gene regulatory networks from time-stamped single cell transcriptional expression profiles. Bioinformatics, 34(2), 258–266.

Pedregosa, F. et al. (2011). Scikit-learn: Machine learning in Python. Journal of Machine Learning Research, 12, 2825–2830.

Robinson, M. D. and Smyth, G. K. (2008). Small-sample estimation of negative binomial dispersion, with applications to SAGE data. Biostatistics, 9(2), 321–332.

Sanchez-Castillo, M. et al. (2018). A bayesian framework for the inference of gene regulatory networks from time and pseudo-time series data. Bioinformatics, 34(6), 964–970.

Specht, A. T. and Li, J. (2017). Leap: constructing gene co-expression networks for single-cell rna-sequencing data using pseudotime ordering. Bioinformatics, 33(5), 764–766.

Tibshirani, R. (1996). Regression shrinkage and selection via the lasso. Journal of the Royal Statistical Society B, 58, 267–288.

Vallejos, C. A. et al. (2015). BASiCS: Bayesian analysis of single-cell sequencing data. PLOS Computational Biology, 11(6), 1–18.

Whitfield, M. et al. (2002). Identification of genes periodically expressed in the human cell cycle and their expression in tumors. Mol Biol Cell, 13(6), 1977–2000.

Zappia, L. et al. (2017). Splatter: simulation of single-cell RNA sequencing data. Genome biology, 18(1).

Zou, H. and Hastie, T. (2005). Regularization and variable selection via the elastic net. Journal of the Royal Statistical Society: Series B (Statistical Methodology), 67(2), 301–320.

